# Erosion of phylogenetic diversity in Neotropical bat assemblages: findings from a whole-ecosystem fragmentation experiment

**DOI:** 10.1101/534057

**Authors:** Sabhrina G. Aninta, Ricardo Rocha, Adrià López-Baucells, Christoph F. J. Meyer

**Author notes:** **Corresponding author**: Sabhrina G. Aninta. Our study highlights the erosion of phylogenetic diversity of bat assemblages associated with habitat fragmentation and degradation in the world’s largest and longest-running whole-ecosystem fragmentation experiment. This work advances our understanding of the effects of habitat modification on bats as key ecosystem service providers in tropical ecosystems.

## Abstract

The traditional focus on taxonomic diversity metrics for investigating species responses to habitat loss and fragmentation has limited our understanding on how biodiversity is impacted by habitat modification. This is particularly true for taxonomic groups such as bats which exhibit species-specific responses. Here, we investigate phylogenetic alpha and beta diversity of Neotropical bat assemblages across two environmental gradients, one in habitat quality and one in habitat amount. We surveyed bats in 39 sites located across a whole-ecosystem fragmentation experiment in the Brazilian Amazon, representing a gradient of habitat quality (interior-edge-matrix, hereafter IEM) in both continuous forest and forest fragments of different sizes (1, 10, and 100 ha; forest size gradient). For each habitat category, we quantified alpha and beta phylogenetic diversity, then used linear models and cluster analysis to explore how forest area and IEM gradient affect phylogenetic diversity. We found that the secondary forest matrix harboured significantly lower total evolutionary history compared to the fragment interiors, especially the 1 ha fragments, containing bat assemblages with more closely related species. Forest fragments ≥ 10 ha had levels of phylogenetic richness similar to continuous forest, suggesting that large fragments retain considerable levels of evolutionary history. The edge and matrix adjacent to large fragments tend to have closely related lineages nonetheless, suggesting phylogenetic homogenization in these IEM gradient categories. Thus, despite the high mobility of bats, fragmentation still induces considerable levels of erosion of phylogenetic diversity, suggesting that the various evolutionary history might not be able to persist in present-day human-modified landscapes.

## Introduction

Humans have fundamentally changed the face of the Earth, with negative side-effects for biodiversity across all major biomes. Tropical forests are among the biomes impacted most heavily given the large footprint of pervasive land use changes which have resulted in widespread landscape fragmentation and loss of habitat for many species (Austin et al. 2017; Barlow et al. 2018). These land use changes have been documented to have detrimental effects on the richness, abundance, and composition of many tropical taxa (Alroy 2017), leading to a pattern of extensive defaunation, with cascading effects on ecosystem functioning (Young et al. 2016).

Idiosyncratic responses of species to fragmentation are ubiquitous, rendering assemblage-level inferences regarding fragmentation effects generally difficult (Ewers and Didham 2006; Fahrig 2017). This is mainly because the treatment of species as equal entities by neglecting their unique evolutionary history, functional roles in the ecosystem, and their association with each other within the community (Pellens and Grandcolas 2016), paints an incomplete picture of the effects of habitat fragmentation. Therefore, recent studies assessing the effect of habitat fragmentation on tropical taxa have started to incorporate evolutionary information using phylogenetic diversity metrics in addition to species richness (Frishkoff et al. 2014; Santos et al. 2014; Cisneros et al. 2015, 2016; Aguirre et al. 2016; Frank et al. 2017). By doing so, these studies were able to uncover patterns previously undetected by studies with a sole focus on the taxonomic dimension of biodiversity. For example, the random pattern of plant species absence across the phylogenetic tree in a fragmented landscape, or phylogenetic over-dispersion, can potentially be linked to low phylogenetic conservatism of key life-history traits (Santos et al. 2014), suggesting that the negative effect of habitat fragmentation depends on the evolutionary history of the taxa in question. However, studies on mobile species such as birds and bats showed non-random patterns of absence across their phylogeny in various types of disturbed habitats (Riedinger et al. 2013; Frishkoff et al. 2014; Frank et al. 2017). This pattern, often referred to as phylogenetic clustering, whereby species within an assemblage are more closely related than expected by chance, indicates a strong effect of habitat filtering. Gaining better insights into the extent to which phylogenetic diversity of assemblages is eroded as a result of habitat fragmentation therefore is critical to improve our general understanding of biodiversity persistence in human-modified landscapes.

Despite their mobility, bats (Chiroptera) are among the many animal groups that are demonstrably affected by habitat loss and fragmentation (Meyer et al. 2016; Alroy 2017). Notwithstanding increased research effort devoted over recent years to better understand how bats respond to habitat fragmentation, studies at the assemblage level, typically comparing species richness, diversity, and assemblage composition between forest fragments and continuous forest, showed inconsistent results and highlighted the need for more research focusing on the functional and phylogenetic biodiversity dimensions (Meyer et al. 2016). Bats are a good model group to study the effect of habitat fragmentation on phylogenetic diversity given their high species richness, functional diversity, and key roles in ecosystem functioning (Kunz et al. 2011). Studies employing a phylogenetic approach to investigate bat responses towards habitat disturbance (Cisneros et al. 2015; Frank et al. 2017; Presley et al. 2018) have been made possible by the availability of phylogenetic trees of all extant bat species (Jones et al. 2002, 2005; Shi and Rabosky 2015) that can be used to calculate phylogenetic diversity metrics.

Of the few studies that have investigated bat phylogenetic diversity in fragmented landscapes, none has assessed responses across the entire gradient in habitat quality typically encountered, formed by the interiors (I) and edges (E) of continuous forest and forest fragments, as well as the intervening matrix (M), or the IEM gradient (Rocha et al. 2017). Explicit consideration of the full IEM gradient, however, is important to better understand the extent of environmental filtering that usually is regarded as the cause of phylogenetic clustering in disturbed habitats (Riedinger et al. 2013; Frank et al. 2017; Presley et al. 2018) as species persistence in fragmented landscapes may be differentially affected by this gradient in habitat quality (Ferreira et al. 2017). The observed phylogenetic richness and structure in each habitat that comprises the IEM gradient can give an indication about the amount of evolutionary history retained by the constituent habitat elements of a fragmented landscape (Cisneros et al. 2015). Moreover, exploring which habitats share similar evolutionary history or harbour lineages that are more closely related compared to other habitats may give insights into the evolution of habitat preferences (Graham and Fine 2008).

To elucidate how habitat fragmentation affects the evolutionary dimension of bat diversity, we investigated the changes in phylogenetic alpha and beta diversity of Amazonian bat assemblages across two environmental gradients, one in habitat quality (IEM gradient) and one in habitat amount (forest size: continuous forest; fragments of 1, 10, 100 ha), in the experimentally fragmented landscape of the Biological Dynamics of Forest Fragments Project (BDFFP), the world’s largest and longest-running experimental study of habitat fragmentation (Haddad et al. 2015). For phylogenetic alpha diversity, we expected assemblages in the secondary forest matrix to retain the least total evolutionary history due to selection of bat lineages that are best adapted to it (Rocha et al. 2018), followed by edges, while forest interiors were predicted to harbour the most total evolutionary history due to greatest resource availability (Ries and Sisk 2004). Accordingly, we predicted phylogenetic diversity to increase with forest size, and phylogenetic clustering to be associated with smaller forest fragments and strongest in the matrix. Finally, we expected that habitat filtering will leave each habitat of the IEM gradient with its own set of unique assemblages so that phylogenetic beta diversity will show that the same category of IEM gradient will contain similar lineages and a similar amount of total evolutionary as a consequence of low phylogenetic turnover.

## Methods

### Study Area

The BDFFP spans ∼1,000 km^2^ and is located approximately 80 km north of Manaus, Brazil (S2°30’, W60°). The area contains different-sized forest fragments separated by 80-650 m from the surrounding continuous forest that serves as experimental control (Laurance et al. 2018). Following abandonment of the cattle pastures which initially surrounded the experimentally isolated fragments after their creation in the early 1980s, *Vismia-* and *Cecropia*-dominated secondary forest developed in the matrix (Mesquita et al. 2015).

The forest at the BDFFP is a typical non-flooded forest of the Amazon basin (De Oliveira and Mori 1999), with approximately 280 species of trees (dbh >10 cm) per hectare (Laurance et al. 2010). The area has a relatively flat topography (80-160 m), with nutrient-poor soils. Rainfall ranges from 1,900 to 3,500 mm annually, with a moderately strong dry season from June to October (Laurance et al. 2018).

### Bat Sampling

Bats were sampled with ground-level mist nets in eight primary forest fragments —three of 1 ha, three of 10 ha and two of 100 ha—and in nine control sites in three areas of continuous forest. Fragments and controls were sampled in the interior, at the edges, and in the secondary forest matrix, resulting in a total of 39 sites (for continuous forest, only three edge and matrix sites were sampled; cf. Rocha et al. 2017). Distances between interior and edge sites of continuous forest and fragments were respectively 1,118 ± 488 and 245 ± 208 m (mean ± SD). Matrix sites were located ca. 100 m away from the border between primary and secondary forest. Each sampling site was visited eight times over a 2-year period (August 2011 – June 2013). See (Rocha et al. 2017) for a more detailed description.

### Phylogenetic Information

Analysis was restricted to the family Phyllostomidae, which can be adequately sampled with ground-level mist nets (Kalko 1998). The phylogenetic information of phyllostomid species present at the BDFFP, hereafter referred to as the local phylogeny, was extracted from the most recent species-level phylogeny of bats (Shi and Rabosky 2015) using R package ‘picante’ (Kembel et al. 2010). We chose this particular tree as it covered more of the phyllostomid species that occur at the BDFFP compared to another frequently used bat phylogenetic tree published by (Jones et al. 2002, 2005). The supertree was downloaded from TreeBASE and pruned to obtain the local phylogeny. The branch lengths of the local phylogeny represent the divergence time of the species in millions of years. As we used distance-based phylogenetic diversity metrics, the phylogenetic pairwise distance matrix was extracted from the local phylogeny using the ‘cophenetic.phylo()’ function from R package ‘ape’ (Paradis et al. 2004).

Species that occur in the study area but were not present in the pruned tree were substituted by their congeners, following (Cisneros et al. 2016). Only for two out of 43 captured phyllostomid species was this the case, i.e. *Artibeus gnomus* and *A. cinereus*, which hence were both represented by their closest congener, *A. glaucus* (Redondo et al. 2008).

### Measuring Phylogenetic Diversity

#### Alpha Diversity

We explored variation in phylogenetic richness and structure within assemblages separately across the IEM gradient of 1, 10, and 100 ha fragments and continuous forest (CF) using Faith’s phylogenetic diversity (Faith 1992) and mean pairwise distance (Clarke and Warwick 1998; Webb 2000), hereafter referred to as PD and MPD respectively. As PD is sensitive to sample size (Vamosi et al. 2009), we rarefied the observed values of PD when comparing different assemblages using the R package ‘BAT’(Cardoso et al. 2015).

To assess the effect of phylogenetic information *per se* on PD, we eliminated any potential effect of species richness by calculating SES_PD_ (standardized effect size of PD) under a ‘richness’ null model (Swenson 2014). Similarly, we quantified phylogenetic structure *per se* by calculating SES_MPD_ (standardized effect size of MPD) using the tip-shuffling null model (Swenson 2014). However, as the local phylogeny consisted of closely related phyllostomids with quite a balanced topology, MPD could underestimate phylogenetic clustering in the terminal part of the phylogeny. We therefore also calculated the mean nearest taxon distance (MNTD) and computed SES_MNTD_ using the tip-shuffling null model as it is more sensitive than MPD for detecting phylogenetic clustering in the terminal part of the local phylogeny (Tucker et al. 2017). For SES_MPD_ and SES_MNTD_, high quantiles (p>0.95) indicate a phylogenetically over-dispersed assemblage whereas low quantiles (p<0.05) indicate a phylogenetically clustered assemblage (Swenson 2014).

#### Beta Diversity

To explore between-assemblage variation in phylogenetic richness and structure in relation to the habitat quality (IEM) and size gradients, we calculated COMDIST and phylogenetic beta diversity (Pβ_total_), respectively. COMDIST is the mean pairwise phylogenetic distance (MPD) between species from different assemblages (Swenson 2011) which could help reveal which habitats along the gradient contain closely related lineages. To calculate COMDIST, we used R package ‘picante’ (Kembel et al. 2010). Phylogenetic beta diversity was partitioned into its richness (Pβ_rich_) and replacement (Pβ_repl_) components to capture, respectively, the difference in shared total branch lengths between assemblages and the uniqueness of each assemblage based on the evolutionary lineages present (Cardoso et al. 2014). To calculate Pβ_total_and its partitions, we used R package ‘BAT’ (Cardoso et al. 2015).

### Phylogenetic alpha and beta diversity across the IEM and forest size gradient

#### Alpha Diversity

We modelled phylogenetic diversity in relation to the IEM and forest size gradient to understand how the two affect the evolutionary dimension of biodiversity. While in principle the six camp locations (Colosso, Porto Alegre, Dimona, Cabo Frio, Florestal and Km 41) should be incorporated in the statistical model as a random effect to account for potential spatial autocorrelation of the response variables (Bolker et al. 2009), we found its effect to be negligible in practice after we fit linear mixed models using the ‘lme4’ R package (Bates et al. 2014) (see Table S1). We therefore dropped the block effect, leaving us with a simple linear model to assess significance of the fixed effects. The linear models were fitted using the R package ‘stats’ (R Core Team 2017), with IEM gradient (interior, edge, or matrix) and forest size (CF, 100 ha, 10 ha, or 1 ha) as explanatory variables, comparing 12 types of assemblages in total. Both additive and interactive effects for IEM gradient and forest size along with the null models were evaluated based on the small sample size version of Akaike’s Information Criterion (AICc) (Burnham and Anderson 2002) as implemented in R package ‘MuMIn’ (Barton 2016). Along with AICc, we also calculated Akaike weights (w_i_) and Akaike differences (Δ_i_) to gauge relative variable importance. Models with Δ_i_ ≤ 2 were considered to have substantial support (Burnham and Anderson 2002). Significant effects were further evaluated via multiple comparison tests with Tukey contrasts (adjusted *P* values reported in Table S3) using the R package ‘multcomp’ (Hothorn et al. 2008).

#### Beta Diversity

To detect any implicit spatial structure in Pβ_total_, Pβ_rich_, Pβ_repl_, and COMDIST, we visualized these metrics through UPGMA clustering (Borcard et al. 2011) using the ‘hclust’ function in R (R Core Team 2017). When applied to Pβ_rich_, UPGMA will cluster assemblages with similar amount of phylogenetic richness whereas Pβ_repl_will cluster assemblages with similar lineages. For COMDIST, UPGMA will cluster closely related assemblages. A Mantel test (Mantel 1967) was conducted using the R package ‘vegan’ (Oksanen et al. 2017) to detect any linear (Pearson correlation) or non-linear (Spearman’s rank correlation) spatial structure in Pβ_total_, Pβ_rich_, Pβ_repl_, and COMDIST.

## Results

### Comparison within assemblages: phylogenetic diversity is lowest in the smallest fragments

Rarefied PD showed a decreasing trend from continuous forest and fragment interiors to forest edges and matrix, with the matrix sites adjacent to the 1 ha fragments harbouring the lowest PD overall (Fig. 1). In agreement with this, the model including IEM gradient and forest size as additive predictors of PD received overwhelming support (*w*_i_ = 0.99) compared to the other candidate models (Table 1). The decrease in PD from interior towards edge and matrix was significant (*P* <0.001 for both), with edges experiencing larger decreases in PD compared to the matrix (Table S2). However, multiple comparison tests did not support any significant differences in PD between edges and matrix, irrespective of forest size (Table S3). Also, there was no significant loss in PD from larger towards smaller fragments except for 1 ha fragments (*P* <0.001, Table S2). Multiple comparison tests confirmed the significantly lower phylogenetic richness in 1 ha fragments relative to larger fragments and continuous forest across the entire IEM gradient (Table S3). According to the local phylogeny, most of the old species lineages, i.e. those with long tree branches, do not appear in the 1 ha fragments (Fig. S1).

**Table 1.**
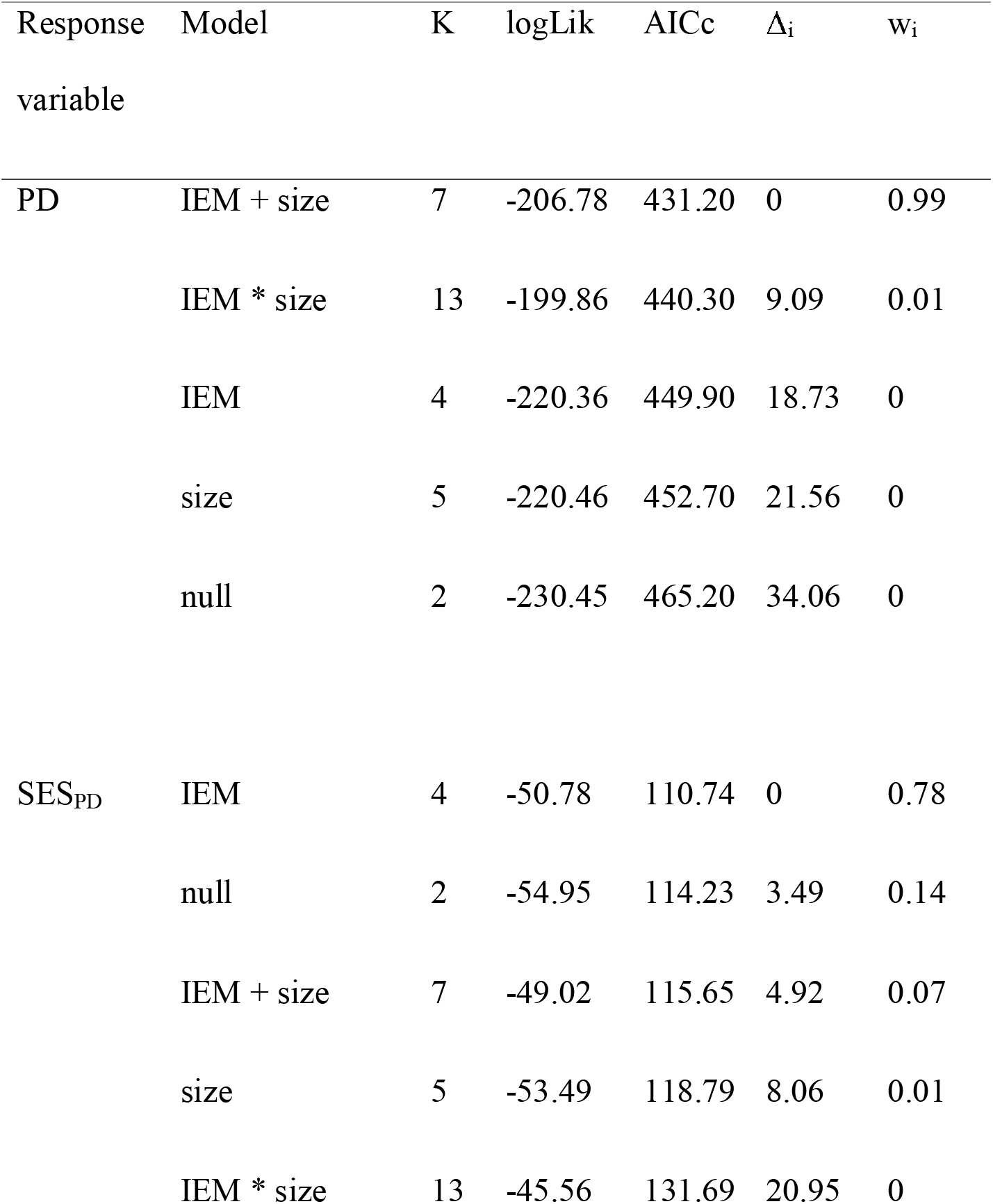
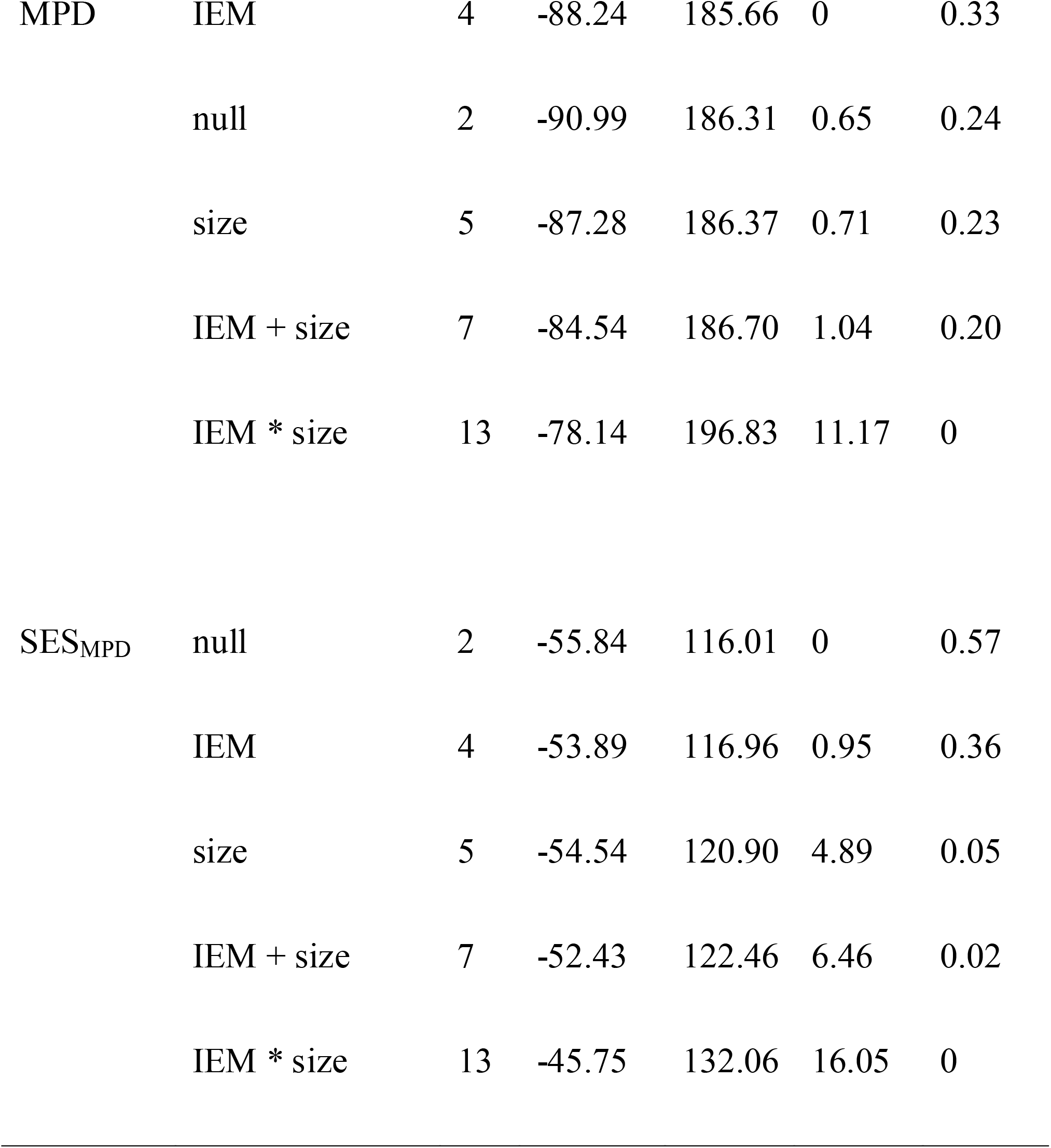
Results of Akaike Information Criterion (AIC)-based model selection assessing support for an effect of habitat quality (IEM) and habitat amount (size) on phylogenetic richness (PD and SES_PD_) and divergence (MPD and SES_MPD_) of bat assemblages at the Biological Dynamics of Forest Fragments Project, Brazil. For each model, the log-likelihood (Log-L), number of estimable parameters (K), sample-size adjusted AIC (AICc), Akaike differences (Δ_i_), and Akaike weights (w_i_) are presented.

**Fig. 1.**
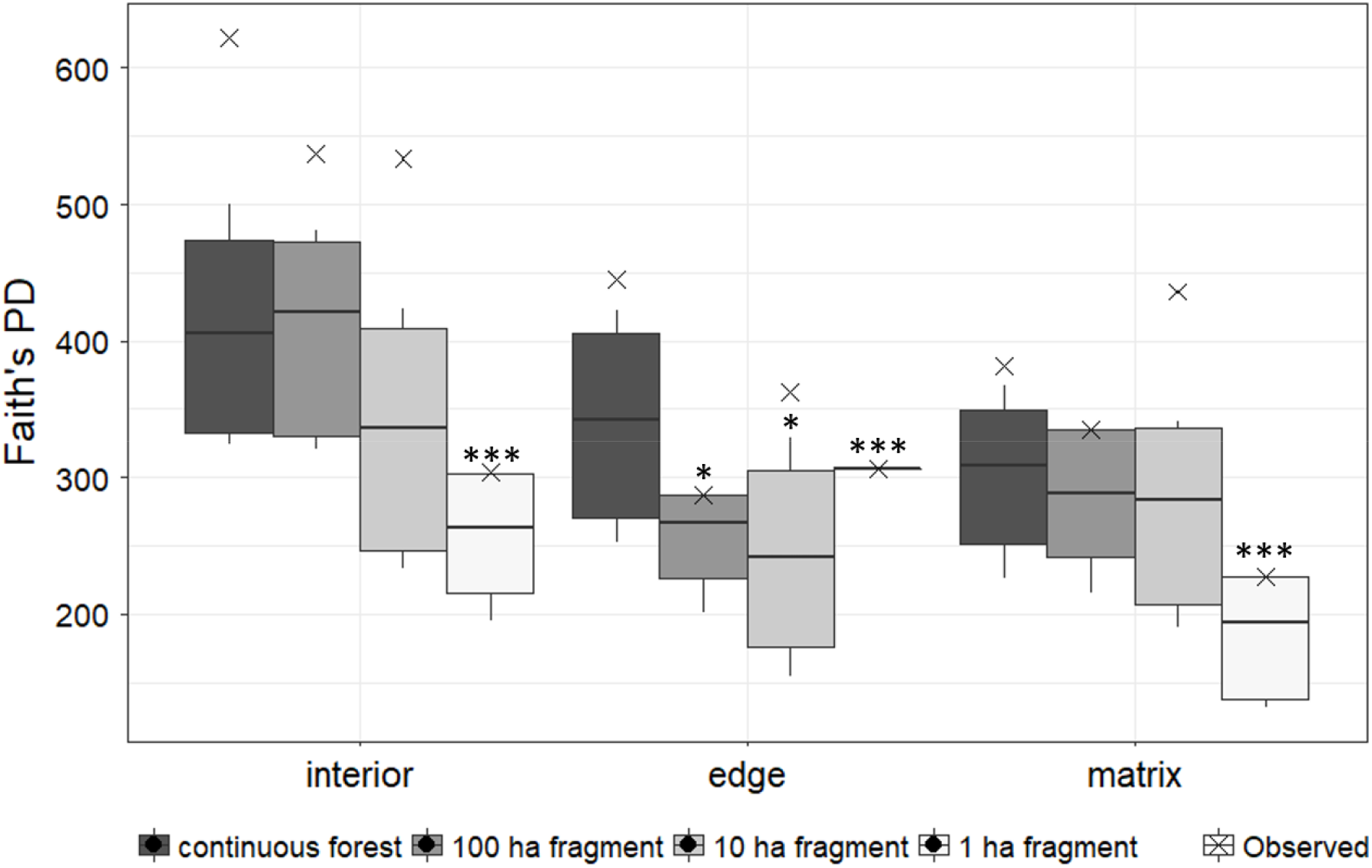
Comparison of rarefied phylogenetic richness of bat assemblages across gradients of habitat quality (IEM) and habitat amount (forest size) at the Biological Dynamics of Forest Fragmentation Project, Brazil. Horizontal bars indicate the median of rarefied values, the mean of observed values for each habitat category is depicted as ‘x’. Asterisks denote significant differences relative to continuous forest interiors (\*\*\**P*<0.001; \*\**P*<0.01; \**P* <0.05).

When the effect of species richness on PD was accounted for (SES_PD_), only the IEM gradient emerged as the best-supported predictor in the model (*w*_i_ = 0.78, Table 1). SES_PD_ for forest fragments showed lower phylogenetic richness than expected given the number of species present, particularly for fragment edges and the matrix (Fig 2A). SES_PD_ was also significantly lower in the matrix than in forest interiors, particularly for the 1 ha fragments (Fig 2A).

**Fig. 2.**
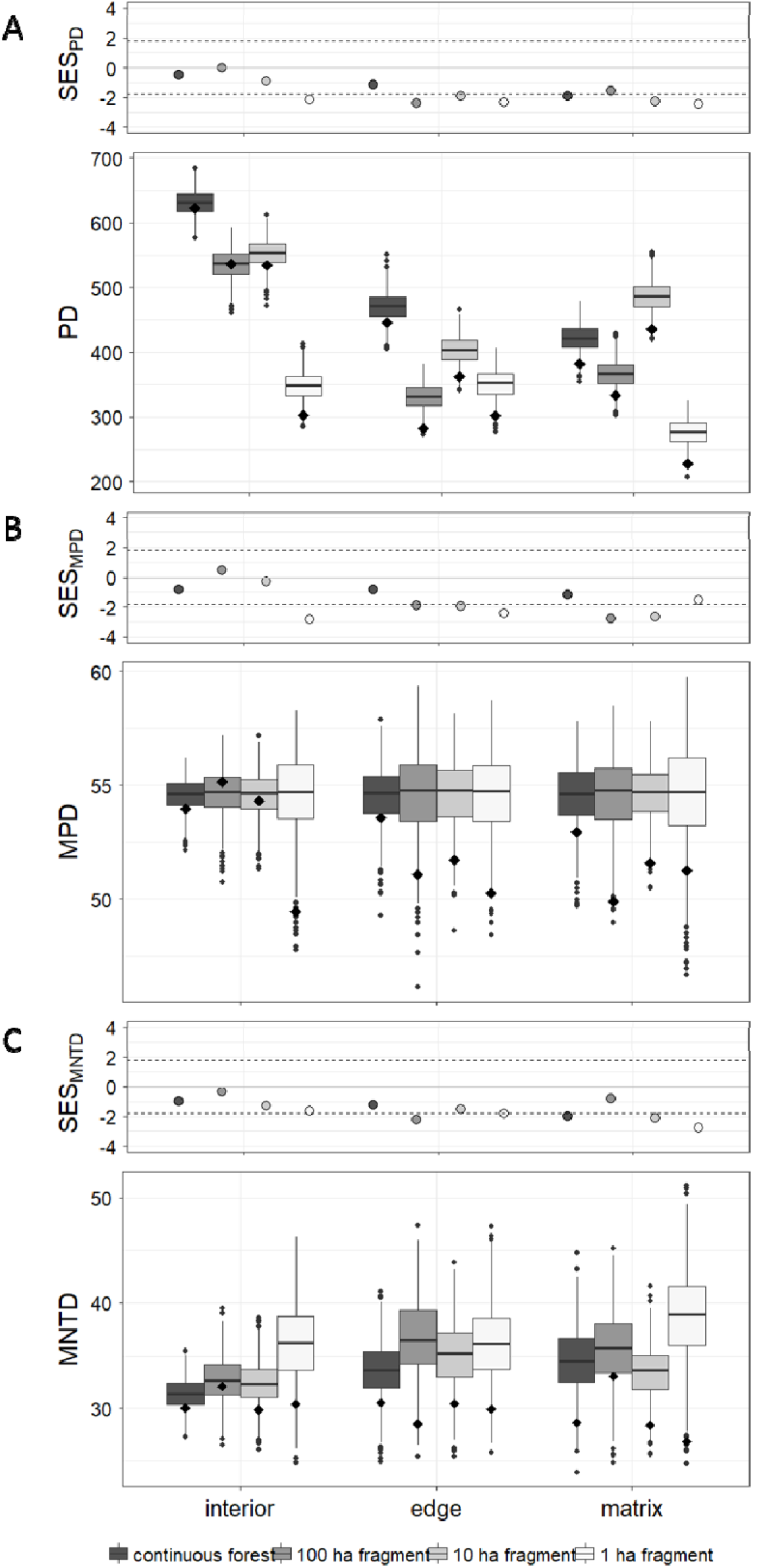
Standardized effect size (dots) of (A) Faith’s phylogenetic diversity (PD), (B) mean pairwise distance (MPD), and (C) mean nearest taxon distance (MNTD) along with 5^th^ and 95^th^ quantiles (dashed lines) of the simulated null communities (box and whisker plots). In the box plots, values of observed PD, MPD, and MNTD overlaid on the simulated null communities are indicated by a black diamond.

In line with PD, observed MPD of phyllostomid assemblages in most habitat types was lower than expected under the simulated null communities, particularly for the interiors of 1 ha fragments, fragment edges, and the matrix around the larger (10 and 100 ha) fragments (Fig. 2B). Only the assemblages in the matrix surrounding 1 ha fragments, although characterized as relatively phylogenetically over-dispersed compared to the other assemblages based on SES_MPD_, showed terminal clustering based on its significantly lower SES_MNTD_ (Fig. 2C). Patterns of MPD were best explained by a model that included only the IEM gradient as explanatory variable (w_i_ = 0.33, Table 1). There was, however, substantial model selection uncertainty, with the null, size, and IEM + size models also being supported as plausible (ΔAIC_c_ <2, Table 1). According to the single variable models IEM gradient and size (Table S2), MPD was significantly reduced in the matrix (*P* = 0.03) and 1 ha fragments (*P* = 0.04), respectively, suggesting that the matrix around 1 ha fragments contains more closely related lineages as indicated by its significantly lower SES_MNTD_. For SES_MPD_, the effect of the IEM gradient was even more tenuous (w_i_ = 0.36) as the null model received the strongest support (w_i_ = 0.57, Table 1).

### Comparison between assemblages: no clear pattern of lineage replacement

Dendrograms based on the total evolutionary history shared between assemblages (Pβ_total_, Pβ_rich_, and Pβ_repl_) were substantially different from the one based on relatedness between lineages within assemblages (COMDIST). UPGMA clustering based on total phylogenetic beta diversity (Pβ_total_) suggested that the interiors of continuous forest and forest fragments harbour similar amounts of phylogenetic richness, except for the 1 ha fragments (Fig 3A), further confirming the phylogenetic erosion of 1 ha fragments. The similarity in total phylogenetic richness between the interiors of continuous forest and larger forest fragments (10 and 100 ha) was maintained after Pβ_total_ was partitioned into Pβ_rich_ and Pβ_repl_. For Pβ_rich_, however, the interior sites of the larger fragments clustered together, unlike Pβ_total_ which grouped the interior of continuous forest closer together with that of 100 ha fragments (Fig 3B). For Pβ_repl_, the interior sites were completely dispersed (Fig 3C), suggesting that the difference in Pβ_total_ compared to Pβ_rich_ is caused by different lineages contained within the interiors. The position of the IEM habitat categories in the Pβ_repl_ dendrogram are unlikely due to their geographic proximity as there was no significant relationship between Pβ_repl_ and geographic distance based on the Mantel test (Pearson’s Mantel statistic r = 0.031, *P* = 0.299, Table S4).

**Fig. 3.**
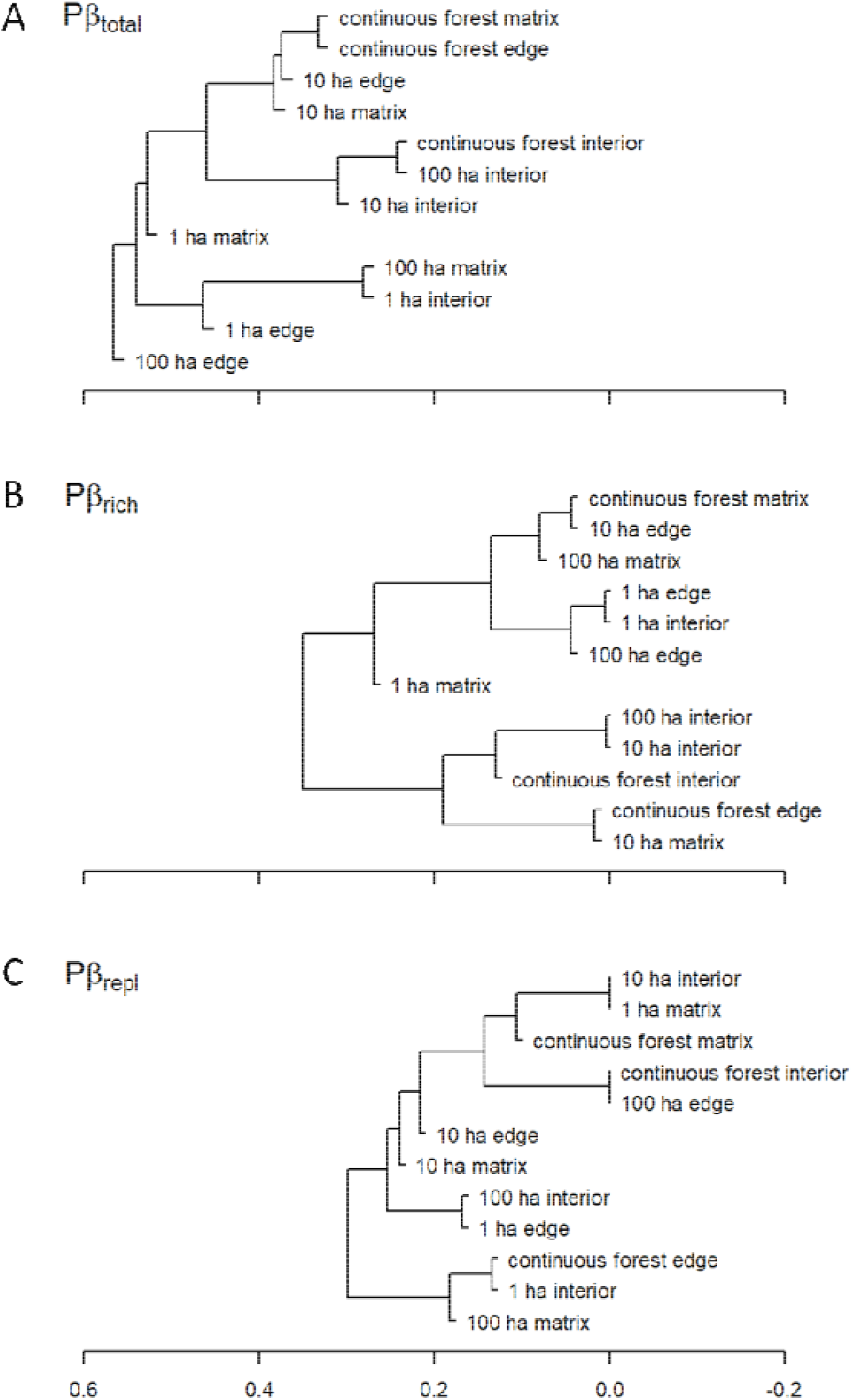
Unweighted Pair Group Method with Arithmetic Mean (UPGMA) clustering of bat total phylogenetic beta richness (Pβ_tot_) (A) in the Biological Dynamics of Forest Fragments Project landscape, partitioned into its richness (Pβ_rich_) (B) and replacement (Pβ_repl_) (C) components.

UPGMA clustering of COMDIST revealed that the investigated assemblages were closely related and were not clustered according to either the same IEM gradient or the same forest size categories (Fig. 4). The sites within the interior of 1 ha fragment are more closely related to the matrix and edges of forest fragments and continuous forest. The position of the interior sites of 1 ha fragments distant from those of larger fragments and continuous forest, underscores the distinct composition of bat lineages in the assemblages of 1 ha fragments.

**Fig. 4.**
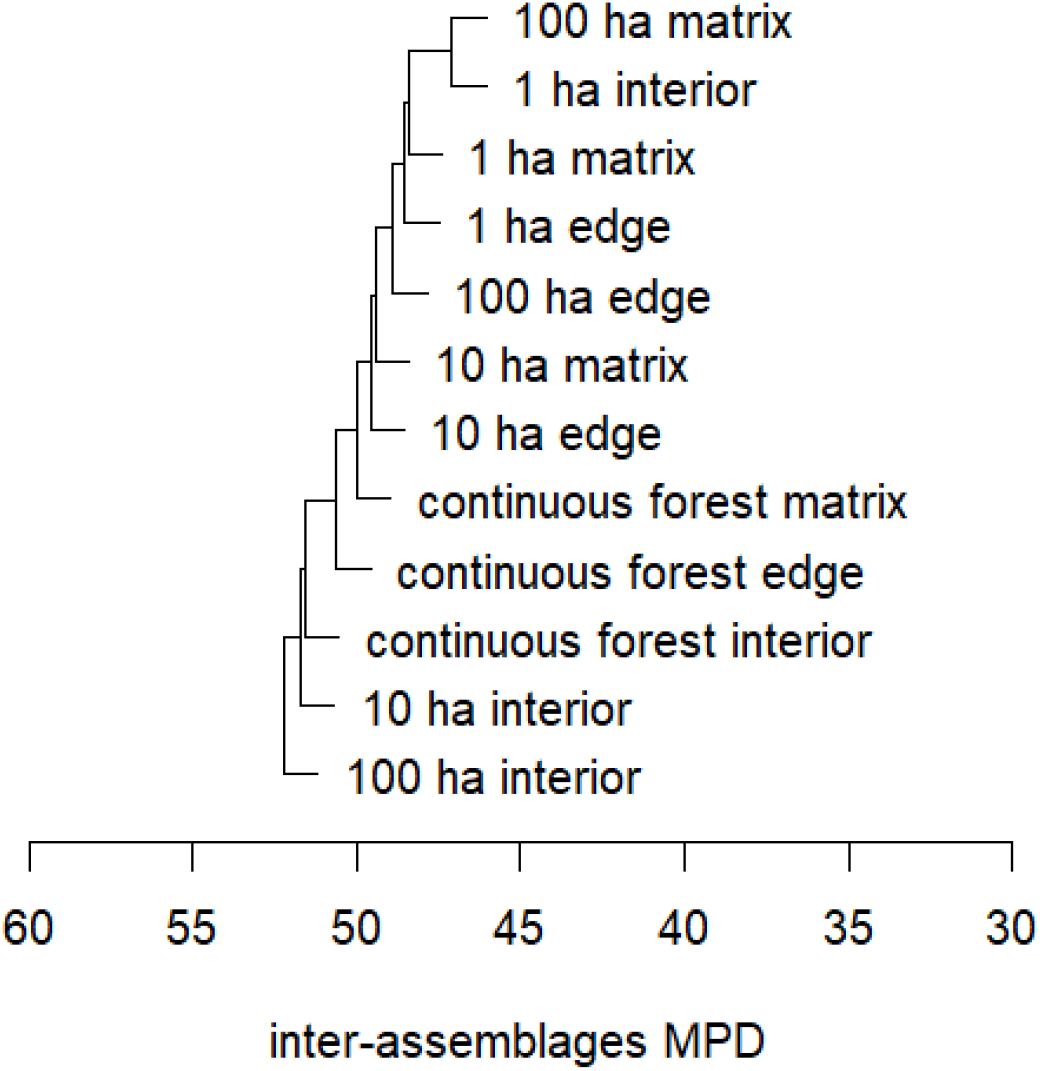
Unweighted Pair Group Method with Arithmetic Mean (UPGMA) clustering of COMDIST metric representing the divergence aspects of phylogenetic beta diversity of bats in the Biological Dynamics of Forest Fragments Project landscape.

When the local phylogeny was annotated with information on species’ presence across the two environmental gradients, different lineages were present within the same IEM category except for the interior of continuous forest and the larger forest fragments (Fig. S1). The interiors of 1 ha fragments were distinct from other interior sites whereas there was no clear pattern of lineage distribution at forest edges and in the matrix adjoining continuous forest and fragments. This is also unlikely due to spatial effects as there was no significant relationship between COMDIST and geographic distance (Pearson’s Mantel statistic r = 0.055, *P* = 0.230, Table S4).

## Discussion

Using metrics of phylogenetic alpha and beta diversity, we showed that there was not only a disproportionate reduction of phylogenetic richness in the smallest forest fragments (1 ha) but also phylogenetic homogenization of assemblages at forest edges and in the matrix as assemblages in these categories of the IEM gradient contained closely related bat lineages. Whereas the two environmental gradients investigated in this study explained quite well the observed variation in total phylogenetic richness (PD), they could not adequately account for how closely related lineages clustered in small forest fragments (MPD). Additionally, the results of the clustering analysis based on phylogenetic beta diversity metrics were counter to our predictions since the resulting dendrograms did not clearly cluster assemblages from the same category of habitat quality and forest size together.

### Environmental filtering is strongest in the matrix surrounding small forest fragments

Across the full disturbance gradient of interior-edge-matrix, phylogenetic richness was curtailed in both edge and matrix habitats, especially in the case of the 1 ha fragments. Thus, habitat fragmentation at the BDFFP does not only reduce bat taxonomic and functional diversity in the small forest fragments, edges, and matrix as we have previously shown (Farneda et al. 2015; Rocha et al. 2017), but also negatively affects the total evolutionary history preserved. However, the magnitude of loss of phylogenetic richness at fragment edges and in the adjacent matrix relative to the interior of continuous forest was only slight, probably due to the low structural contrast between matrix (tall secondary forest) and edges during the sampling period (Rocha et al. 2017). In accordance with other studies comparing phylogenetic richness between various human-modified habitats of different levels of forest/vegetation complexity (Riedinger et al. 2013; Frishkoff et al. 2014; Edwards et al. 2015, 2017; Frank et al. 2017; Martins et al. 2017; Pereira et al. 2018), the observed decrease in total evolutionary history in a structurally less complex habitat such as the secondary forest matrix is not surprising. Phylogenetic diversity is often associated with canopy cover (Frank et al. 2017; Martins et al. 2017), which decreased along our IEM gradient (cf Table S2, Rocha et al. 2017). This phenomenon is often attributed to environmental filtering, whereby species with particular characteristics are not well adapted to and thus unable to persist in all parts of an environmental gradient (Cisneros et al. 2016; Frank et al. 2017).

Through the use of SES_MPD_ and SES_MNTD_, we further detected that edge and matrix assemblages were phylogenetically clustered, whereby the matrix of 1 ha fragments was characterized by especially closely related lineages, pointing towards stronger environmental filtering compared to the edge. Matrix sites surrounding 1 ha fragments are impoverished with respect to lineages with long evolutionary history, i.e. *Lampronycteris brachyotis* and several *Micronycteris* species (Fig S1), and contain lineages that are phylogenetically clustered in the terminal branches. Nevertheless, it is important to note that matrix sites adjacent to the larger fragments (10 ha and 100 ha) did not differ much from their edges in terms of phylogenetic richness and structure, indicating that the habitat quality gradient investigated in this study represents a strong environmental filter in both large (≥ 10 ha) and small (1 ha) forest fragments.

Our model selection results, however, did not provide strong support for the environmental gradient investigated in this study to explain the observed phylogenetic clustering, which was weak as the values of SES_MPD_ and and SES_MNTD_ did not deviate much from the 5^th^ and 95^th^ quantiles of the null model. This is potentially due to the influence of landscape-level attributes which encompass wider environmental gradients (Rocha et al. 2017; Tinoco et al. 2018) and are responsible for the ability of the large fragments to retain more distantly related lineages than the small forest fragments. The low structural contrast between secondary regrowth in the matrix and the forest interiors during the sampling period could also attenuate any strong phylogenetic structure of bat assemblages along the interior-edge-matrix gradient; (Patrick and Stevens 2016) for instance found that environmental variables have a weaker effect upon phylogenetic structure when assemblages experience less harsh environmental conditions. Additionally, the environmental filtering that is responsible for phylogenetic clustering could possibly result from non-linear responses (Smith and Lundholm 2010; Stegen and Hurlbert 2011) that we were unable to detected using linear models. Stronger support of an effect of the IEM gradient in explaining phylogenetic structure may have been received if we had considered abundance-based metrics and quantitative variables such as seasonal temperature (Stevens and Gavilanez 2015) or elevation (Cisneros et al. 2014). Nonetheless, our findings still confirm our expectations that the low resource availability in the matrix apparently selects for frugivorous bat lineages that have traits favourable by the existing environmental filter (Farneda et al. 2015).

There is a possibility that the phylogenetic clustering documented in small fragments also reflects biotic filtering through competitive exclusion of species with low competitive ability in addition to abiotic filtering through selection by the IEM gradient (Mayfield and Levine 2010). However, we consider this unlikely to be the case for phyllostomid bats which are known to have a high diversification rate compared to other bat families (Monteiro and Nogueira 2011; Santana et al. 2012). The wide array of feeding strategies exhibited by phyllostomids likely eliminates the need for strong competition between species as pre-requisite of inferring biotic filtering. For bats at the BDFFP in particular, traits associated with fragmentation sensitivity such as trophic level, body mass, and wing morphology were phylogenetically conserved (Farneda et al. 2015) so that the observed clustering could be related to environmental filtering.

By investigating two measures of phylogenetic alpha diversity simultaneously (PD and MPD), we showed that assemblages that have a relatively lower amount of phylogenetic richness (PD) compared to those of the interior of continuous forest also are made up of more closely related lineages as shown by the relative increase in phylogenetic clustering (MPD). This is consistent with Frank et al (2017) and Riedinger et al (2013) who pointed out the tendency of closely-related bat species to respond similarly to habitat disturbance due to the similarity in traits that are evolutionarily conserved.

### Variation in phylogenetic diversity between assemblages is not related to habitat quality

Phylogenetic beta diversity metrics further corroborate the weak effect of habitat quality on between-assemblage differences in phylogenetic richness and lineage composition. There was no clear-cut pattern with regard to the relatedness and phylogenetic richness of bat assemblages in relation to IEM and forest size gradients. Except for 1 ha fragments, the interiors of the forest fragments and continuous forest contain a large number of distantly related species and harbour a similar amount of total evolutionary history, but not the same bat lineages. Larger forest fragments (≥10 ha) may therefore act as important repositories for preserving the total evolutionary history of bat assemblages in fragmented landscapes. In contrast, the marked changes in phylogenetic richness and structure of 1 ha fragment interiors and similarity to their edges in terms of preserved evolutionary history, suggests that patterns of phylogenetic diversity are fundamentally driven by the pervasive negative edge effects that commonly plague fragments of this size (Santos et al. 2010; Laurance et al. 2018). We probably did not detect any spatially implicit structure between different habitat categories of the IEM gradient due to the predominant effect of dispersal ability in the absence of contrasting environmental gradients (Moreno and Halffter 2001).

The distribution of edge and matrix sites associated with the larger forest fragments and continuous forest across the dendrograms are possibly the result of phylogenetic homogenization. Assemblages of these two categories of the IEM gradient did not differ markedly in phylogenetic richness and structure according to the phylogenetic alpha diversity metrics, and UPGMA clustering further confirmed that they are closely related to each other (Fig. 4). Thus, despite the ability of large forest fragments to retain amounts of total evolutionary history similar to the interior of continuous forest, the level of forest degradation at the edges and in the matrix still has a negative effect on the phylogenetic diversity of bats at the BDFFP.

### New insights gained from a focus on the phylogenetic biodiversity dimension

A focus on the phylogenetic dimension of biodiversity has allowed us to complement our previous analyses that approached bat responses to landscape fragmentation at the BDFFP from taxonomic and functional perspectives (Rocha et al. 2017; Farneda et al. 2018). The reduction of total evolutionary history in the smallest fragments, particularly in the surrounding matrix, was apparently selective and resulted in assemblages comprised of closely related bat lineages with similar traits due to environmental filtering. This adds more significance to the term “biodiversity loss” in fragmented landscapes as losing a lineage of species usually means losing a unique evolutionary history which may be irreplaceable for a particular habitat and disrupt ecosystem processes therein. The non-significant decrease of phylogenetic richness in larger fragments (10 ha and 100 ha) compared to continuous forest indicates that fragments of that size still serve as important refuges for the overall lineage pool in the BDFFP landscape. Although the habitat quality and size gradients investigated in this study could not entirely explain the pattern of phylogenetic beta diversity, cluster analysis showed their constituent categories to harbour distinct assemblages that are probably shaped by a specific assembly mechanism.

Any environmental filtering detected in this study, however, may not be generalizable to other fragmented landscapes. The BDFFP as an experimentally controlled fragmented landscape is a best-case scenario as levels of disturbance are reduced in relation to most real-world landscapes (Laurance et al. 2018). Additionally, forest fragments are surrounded by a benign matrix dominated by forest regrowth (Mesquita et al. 2015), and which is largely free from human disturbances which could interact additively or synergistically with fragmentation (Laurance et al. 2018). The matrix was dominated by tall secondary forest regrowth at the time of study (Rocha et al. 2017), resulting in a rather homogenous makeup of the overall landscape which could promote random placement in bat assemblages (Morlon et al. 2011). Nonetheless, the most challenging part in investigating environmental filtering using phylogenetic diversity metrics is deciding which aspect of the environment actually matters for the taxa in question and on which ecological and evolutionary scale to better understand the evolutionary consequences of habitat fragmentation.

Phylogenetic diversity summarizes species characteristics that are not covered by taxon and functional diversity (Flynn et al. 2011; Cisneros et al. 2014), especially the ecological redundancies and complementarity of organisms (Cadotte et al. 2013). Our choice of incidence-based metrics, however, was limited to assess among-species variation and could not account for within-species variation that is better captured by abundance-based metrics. Abundance-based phylogenetic diversity metrics are more sensitive to environmental gradients (Stevens and Gavilanez 2015) and could correct for stochasticity in the detection of rare taxa or sample effects that have been observed for various taxa at BDFFP (Laurance et al. 2018).

Our study adds to a growing body of evidence suggesting biotic homogenization in human-modified landscapes is widespread not only with regard to the taxonomic facet of biodiversity but also its evolutionary dimension (La Sorte et al. 2018; Park and Razafindratsima 2019). Despite the limitation of our methods, we showed fragmentation to result in the loss of species lineages and phylogenetic homogenization of assemblages in degraded edge and matrix habitats. This could further affect ecosystem stability as communities where species are evenly and distantly related to one another are more stable compared to communities where phylogenetic relationships are more clumped (Cadotte et al. 2012). The conservation of large forest fragments and the improvement of habitat quality thus should be priority in managing fragmented forest landscapes to conserve species with longer evolutionary history.

## Acknowledgements

We are in debt to the many volunteers, students and field assistants that helped us during fieldwork and the coordination team of the BDFFP. We also thank Paulo E.D. Bobrowiec for logistic support and Dirk Metzler for his feedback in the statistical analysis.

Author Contributions: RR and AL conducted the fieldwork, SGA and CFJM formulated the idea, SGA performed the analysis, and SGA, RR, AL, and CFJM wrote the manuscript

## References

Aguirre LF, Montano-Centellas FA, Gavilanez MM, Stevens RD (2016) Taxonomic and Phylogenetic Determinants of Functional Composition of Bolivian Bat Assemblages. PLoS One 11:e0158170. doi:10.1371/journal.pone.0158170

Alroy J (2017) Effects of habitat disturbance on tropical forest biodiversity. PNAS 114:6056–6061. doi:10.1073/pnas.1611855114

Austin KG, González-Roglich M, Schaffer-Smith D, et al (2017) Trends in size of tropical deforestation events signal increasing dominance of industrial-scale drivers. Environ Res Lett 12:79601. doi:10.1088/1748-9326/aa7760

Barlow J, Franga F, Gardner TA, et al (2018) The future of hyperdiverse tropical ecosystems. Nature 559:517. doi:10.1038/s41586-018-0301-1

Barton K (2016) MuMIn: Multi-Model Inference.

Bates D, Mächler M, Bolker B, Walker S (2014) Fitting linear mixed-effects models using lme4. arXiv preprint arXiv:14065823

Bolker BM, Brooks ME, Clark CJ, et al (2009) Generalized linear mixed models: a practical guide for ecology and evolution. Trends Ecol Evol 24:127–135. doi:10.1016/j.tree.2008.10.008

Borcard D, Gillet F, Legendre P (2011) Numerical Ecology with R. Springer New York, New York, NY

Burnham KP, Anderson DR (2002) Model selection and multimodel inference: a practical information-theoretic approach, 2nd ed. Springer, New York

Cadotte MW, Albert CH, Walker SC (2013) The ecology of differences: assessing community assembly with trait and evolutionary distances. Ecol Lett 16:1234–1244. doi:10.1111/ele.12161

Cadotte MW, Dinnage R, Tilman D (2012) Phylogenetic diversity promotes ecosystem stability. Ecology 93:S223–S233

Cardoso P, Rigal F, Carvalho JC (2015) BAT - Biodiversity Assessment Tools, an R package for the measurement and estimation of alpha and beta taxon, phylogenetic and functional diversity. Methods in Ecology and Evolution 6:232–236. doi:10.1111/2041-210X.12310

Cardoso P, Rigal F, Carvalho JC, et al (2014) Partitioning taxon, phylogenetic and functional beta diversity into replacement and richness difference components. J Biogeogr 41:749–761. doi:10.1111/jbi.12239

Cisneros LM, Burgio KR, Dreiss LM, et al (2014) Multiple dimensions of bat biodiversity along an extensive tropical elevational gradient. J Anim Ecol 83:1124–1136. doi:10.1111/1365-2656.12201

Cisneros LM, Fagan ME, Willig MR (2015) Effects of human-modified landscapes on taxonomic, functional and phylogenetic dimensions of bat biodiversity. Divers Distrib 21:523–533. doi:10.1111/ddi.12277

Cisneros LM, Fagan ME, Willig MR (2016) Environmental and spatial drivers of taxonomic, functional, and phylogenetic characteristics of bat communities in human-modified landscapes. PeerJ 4:e2551. doi:10.7717/peerj.2551

Clarke KR, Warwick RM (1998) A taxonomic distinctness index and its statistical properties. Journal of Applied Ecology 35:523–531

De Oliveira AA, Mori SA (1999) A central Amazonian terra firme forest. I. High tree species richness on poor soils. Biodivers Conserv 8:1219–1244

Edwards DP, Gilroy JJ, Thomas GH, et al (2015) Land-Sparing Agriculture Best Protects Avian Phylogenetic Diversity. Current Biology 25:2384–2391. doi:10.1016/j.cub.2015.07.063

Edwards DP, Massam MR, Haugaasen T, Gilroy JJ (2017) Tropical secondary forest regeneration conserves high levels of avian phylogenetic diversity. Biological Conservation 209:432–439. doi:10.1016/j.biocon.2017.03.006

Ewers RM, Didham RK (2006) Confounding factors in the detection of species responses to habitat fragmentation. Biol Rev 81:117. doi:10.1017/S1464793105006949

Fahrig L (2017) Ecological responses to habitat fragmentation per se. Annu Rev Ecol Evol Syst 48:1–23. doi:10.1146/annurev-ecolsys-110316-022612

Faith DP (1992) Conservation evaluation and phylogenetic diversity. Biological Conservation 61:1–10

Farneda FZ, Rocha R, López-Baucells A, et al (2015) Trait-related responses to habitat fragmentation in Amazonian bats. J Appl Ecol 52:1381–1391. doi:10.1111/13652664.12490

Farneda FZ, Rocha R, López-Baucells A, et al (2018) Functional recovery of Amazonian bat assemblages following secondary forest succession. Biol Conserv 218:192–199. doi:10.1016/j.biocon.2017.12.036

Ferreira DF, Rocha R, López-Baucells A, et al (2017) Season-modulated responses of Neotropical bats to forest fragmentation. Ecol Evol 7:4059–4071. doi:10.1002/ece3.3005

Flynn DF, Mirotchnick N, Jain M, et al (2011) Functional and phylogenetic diversity as predictors of biodiversity-ecosystem-function relationships. Ecology 92:1573–1581

Frank HK, Frishkoff LO, Mendenhall CD, et al (2017) Phylogeny, Traits, and Biodiversity of a Neotropical Bat Assemblage: Close Relatives Show Similar Responses to Local Deforestation. Am Nat 190:0–0. doi:10.1086/692534

Frishkoff LO, Karp DS, M’Gonigle LK, et al (2014) Loss of avian phylogenetic diversity in neotropical agricultural systems. Science 345:1343–1346

Graham CH, Fine PVA (2008) Phylogenetic beta diversity: linking ecological and evolutionary processes across space in time. Ecol Lett 11:1265–1277. doi:10.1111/j.1461-0248.2008.01256.x

Haddad NM, Brudvig LA, Clobert J, et al (2015) Habitat fragmentation and its lasting impact on Earth’s ecosystems. Science Advances 1:e1500052–e1500052. doi:10.1126/sciadv.1500052

Hothorn T, Bretz F, Westfall P (2008) Simultaneous inference in general parametric models. Biom J 50:346–363

Jones KE, Bininda-Emonds ORP, Gittleman JL (2005) Bats, clocks, and rocks: diversification patterns in Chiroptera. Evolution 59:2243. doi:10.1554/04-635.1

Jones KE, Purvis A, MacLarnon A, et al (2002) A phylogenetic supertree of the bats (Mammalia: Chiroptera). Biol Rev 77:223–259

Kalko EKV (1998) Organisation and diversity of tropical bat communities through space and time. Zoology 101:281–297

Kembel SW, Cowan PD, Helmus MR, et al (2010) Picante: R tools for integrating phylogenies and ecology. Bioinformatics 26:1463–1464. doi:10.1093/bioinformatics/btq166

Kunz TH, Braun de Torrez E, Bauer D, et al (2011) Ecosystem services provided by bats. Annals of the New York Academy of Sciences 1223:1–38. doi:10.1111/j.1749-6632.2011.06004.x

La Sorte FA, Lepczyk CA, Aronson MFJ, et al (2018) The phylogenetic and functional diversity of regional breeding bird assemblages is reduced and constricted through urbanization. Divers Distrib 24:928–938. doi:10.1111/ddi.12738

Laurance SGW, Laurance WF, Andrade A, et al (2010) Influence of soils and topography on Amazonian tree diversity: a landscape-scale study. J Veg Sci 21:96–106. doi:10.1111/j.1654-1103.2009.01122.x

Laurance WF, Camargo JLC, Fearnside PM, et al (2018) An Amazonian rainforest and its fragments as a laboratory of global change. Biol Rev 93:223–247. doi:10.1111/brv.12343

Mantel N (1967) The detection of disease clustering and a generalized regression approach. Cancer Res 27:209–220

Martins ACM, Willig MR, Presley SJ, Marinho-Filho J (2017) Effects of forest height and vertical complexity on abundance and biodiversity of bats in Amazonia. For Ecol Manage 391:427–435. doi:10.1016/j.foreco.2017.02.039

Mayfield MM, Levine JM (2010) Opposing effects of competitive exclusion on the phylogenetic structure of communities: Phylogeny and coexistence. Ecol Lett 13:1085–1093. doi:10.1111/j.1461-0248.2010.01509.x

Mesquita R de CG, Massoca PE dos S, Jakovac CC, et al (2015) Amazon Rain Forest Succession: Stochasticity or Land-Use Legacy? BioScience 65:849–861. doi:10.1093/biosci/biv108

Meyer CFJ, Struebig MJ, Willig MR (2016) Responses of Tropical Bats to Habitat Fragmentation, Logging, and Deforestation. In: Voigt CC, Kingston T (eds) Bats in the Anthropocene: Conservation of Bats in a Changing World. Springer International Publishing, Cham, pp 63–103

Monteiro LR, Nogueira MR (2011) Evolutionary patterns and processes in the radiation of phyllostomid bats. BMC Evol Biol 11:137

Moreno CE, Halffter G (2001) Spatial and temporal analysis of a, ß and y diversities of bats in a fragmented landscape. Biodivers Conserv 10:367–382

Morlon H, Schwilk DW, Bryant JA, et al (2011) Spatial patterns of phylogenetic diversity. Ecol Lett 14:141–149. doi:10.1111/j.1461-0248.2010.01563.x

Oksanen J, Blanchet FG, Friendly M, et al (2017) vegan: Community Ecology Packag

Paradis E, Claude J, Strimmer K (2004) APE: Analyses of Phylogenetics and Evolution in R l nguage. Bioinformatics 20:289–290. doi:10.1093/bioinformatics/btg412

Park DS, Razafindratsima OH (2019) Anthropogenic threats can have cascading homogenizing effects on the phylogenetic and functional diversity of tropical ecosystems. Ecography 42:148–161. doi:10.1111/ecog.03825

Patrick LE, Stevens RD (2016) Phylogenetic community structure of North American desert bats: influence of environment at multiple spatial and taxonomic scales. J Anim Ecol 85:1118–1130. doi:10.1111/1365-2656.12529

Pellens R, Grandcolas P (2016) Phylogenetics and Conservation Biology: Drawing a Path into the Diversity of Life. In: Biodiversity Conservation and Phylogenetic Systematics: Preserving our evolutionary heritage in an extinction crisis. Springer Open, pp 1–15

Pereira MJR, Fonseca C, Aguiar LMS (2018) Loss of multiple dimensions of bat diversity under land-use intensification in the Brazilian Cerrado. HYSTRIX 29:25–32. doi:10.4404/hystrix-00020-2017

Presley SJ, Cisneros LM, Higgins CL, et al (2018) Phylogenetic and functional underdispersion in Neotropical phyllostomid bat communities. Biotropica 50:135145. doi:10.1111/btp.12501

R Core Team (2017) R: A Language and Environment for Statistical Computing. R Foundation for Statistical Computing, Vienna, Austria

Redondo RAF, Brina LPS, Silva RF, et al (2008) Molecular systematics of the genus Artibeus (Chiroptera: Phyllostomidae). Mol Phylogenet Evol 49:44–58. doi:10.1016/j.ympev.2008.07.001

Riedinger V, Müller J, Stadler J, et al (2013) Assemblages of bats are phylogenetically clustered on a regional scale. Basic Appl Ecol 14:74–80. doi:10.1016/j.baae.2012.11.006

Ries L, Sisk TD (2004) A predictive model of edge effects. Ecology 85:2917–2926

Rocha R, López-Baucells A, Farneda FZ, et al (2017) Consequences of a large-scale fragmentation experiment for Neotropical bats: disentangling the relative importance of local and landscape-scale effects. Landscape Ecol 32:31–45. doi:10.1007/s10980-016-0425-3

Rocha R, Ovaskainen O, López-Baucells A, et al (2018) Secondary forest regeneration benefits old-growth specialist bats in a fragmented tropical landscape. Sci Rep 8:3819. doi:10.1038/s41598-018-21999-2

Santana SE, Grosse IR, Dumont ER (2012) Dietary hardness, loading behavior, and the evolution of skull form in bats. Evolution 66:2587–2598. doi:10.1111/j.1558-5646.2012.01615.x

Santos BA, Arroyo-Rodríguez V, Moreno CE, Tabarelli M (2010) Edge-Related Loss of Tree Phylogenetic Diversity in the Severely Fragmented Brazilian Atlantic Forest. PLoS ONE 5:e12625. doi:10.1371/journal.pone.0012625

Santos BA, Tabarelli M, Melo FPL, et al (2014) Phylogenetic Impoverishment of Amazonian Tree Communities in an Experimentally Fragmented Forest Landscape. PLoS ONE 9:e113109. doi:10.1371/journal.pone.0113109

Shi JJ, Rabosky DL (2015) Speciation dynamics during the global radiation of extant bats. Evolution 69:1528–1545. doi:10.1111/evo.12681

Smith TW, Lundholm JT (2010) Variation partitioning as a tool to distinguish between niche and neutral processes. Ecography 33:648–655. doi:10.1111/j.1600-0587.2009.06105.x

Stegen JC, Hurlbert AH (2011) Inferring Ecological Processes from Taxonomic, Phylogenetic and Functional Trait β-Diversity. PLoS ONE 6:e20906. doi:10.1371/journal.pone.0020906

Stevens RD, Gavilanez MM (2015) Dimensionality of community structure: phylogenetic, morphological and functional perspectives along biodiversity and environmental gradients. Ecography 38:861–875. doi:10.1111/ecog.00847

Swenson NG (2014) Functional and Phylogenetic Ecology in R. Springer New York, New York, NY

Swenson NG (2011) Phylogenetic Beta Diversity Metrics, Trait Evolution and Inferring the Functional Beta Diversity of Communities. PLoS ONE 6:e21264. doi:10.1371/journal.pone.0021264

Tinoco BA, Santillân VE, Graham CH (2018) Land use change has stronger effects on functional diversity than taxonomic diversity in tropical Andean hummingbirds. Ecol Evol 8:3478–3490. doi:10.1002/ece3.3813

Tucker CM, Cadotte MW, Carvalho SB, et al (2017) A guide to phylogenetic metrics for conservation, community ecology and macroecology: A guide to phylogenetic metrics for ecology. Biol Rev 92:698–715. doi:10.1111/brv.12252

Vamosi SM, Heard SB, Vamosi JC, Webb CO (2009) Emerging patterns in the comparative analysis of phylogenetic community structure. Mol Ecol 18:572–592. doi:10.1111/j.1365-294X.2008.04001.x

Webb CO (2000) Exploring the phylogenetic structure of ecological communities: an example for rain forest trees. Am Nat 156:145–155

Young HS, McCauley DJ, Galetti M, Dirzo R (2016) Patterns, Causes, and Consequences of Anthropocene Defaunation. Annu Rev Ecol Evol Syst 47:333–358. doi:10.1146/annurev-ecolsys-112414-054142

