## Supplementary material for "Erosion of phylogenetic diversity in Neotropical bat assemblages: findings from a whole-ecosystem fragmentation experiment": Electronic Supplementary Materials (from another version of this preprint: version 1, 2019-01-29)

**Corresponding author**: Sabhrina G. Aninta; **E-mail**: sagitaninta[at]gmail.com

### This supplementary material contains:

**Table S1.** Random effects and residuals of the candidate models assessing the effect of Interior–Edge–Matrix gradient and fragment size on phylogenetic richness and structure.

**Table S2.** Parameter estimates of the best-fit candidate models ($\Delta_i$ <2) explaining the effects of Interior–Edge–Matrix gradient of continuous forest and different-sized fragments on phylogenetic richness and divergence.

**Table S3.** Multiple pairwise comparisons of linear models testing for differences in phylogenetic richness across the Interior–Edge–Matrix gradient of continuous forest and different-sized fragments.

**Table S4.** Mantel tests between beta diversity metrics and geographic distance.

**Fig. S1** Phylogenetic relationship of phyllostomid bats and their occurrence in continuous forest and 100 ha, 10 ha and 1 ha fragments across a gradient of interior, edge, and matrix sites.

**Table S1**. Random effects and residuals of the candidate models assessing the effect of IEM gradient (Gradient) and fragment size (Size) on phylogenetic richness (PD and $SES_{PD}$) and structure (MPD and $SES_{MPD}$) fitted with linear mixed effects models incorporating sampling locations as random effects.

| Response variable | Model | Random Effects | | Residual | | Deviance |
|---|---|---|---|---|---|---|
| | | Variance | Std. Dev | Variance | Std. Dev | |
| **PD** | (1 \| Location) | 959.8 | 30.98 | 7023.5 | 83.81 | 459.8 |
| | Gradient + (1 \| Location) | **$6.23 \cdot 10^{-12}$** | **$2.50 \cdot 10^{-6}$** | 4735.00 | 68.81 | 440.7 |
| | Size + (1 \| Location) | **$4.01 \cdot 10^{-12}$** | **$2 \cdot 10^{-6}$** | 4758.00 | 68.98 | 440.9 |
| | Gradient + Size + (1 \| Location) | 50.81 | 7.13 | 2309.46 | 48.06 | 413.5 |
| | Gradient * Size + (1 \| Location) | 194.80 | 13.96 | 1468.60 | 38.32 | 398.7 |
| **$SES_{PD}$** | (1 \| Location) | 0.01 | 0.11 | 0.97 | 0.98 | 109.9 |
| | Gradient + (1 \| Location) | 0.01 | 0.09 | 0.78 | 0.89 | 101.6 |
| | Size + (1 \| Location) | $4.51 \cdot 10^{-19}$ | $2.13 \cdot 10^{-8}$ | 0.91 | 0.95 | 107.0 |
| | Gradient + Size + (1 \| Location) | $2.40 \cdot 10^{-16}$ | $1.55 \cdot 10^{-8}$ | 0.72 | 0.85 | 98.0 |
| | Gradient * Size + (1 \| Location) | 0.01 | 0.12 | 0.59 | 0.77 | 91.1 |
| **MPD** | (1 \| Location) | 0.06 | 0.25 | 6.16 | 2.50 | 182.0 |
| | Gradient + (1 \| Location) | $5.86 \cdot 10^{-18}$ | $2.42 \cdot 10^{-9}$ | 5.41 | 2.33 | 176.5 |
| | Size + (1 \| Location) | 0 | 0 | 5.14 | 2.27 | 174.6 |
| | Gradient + Size (1 \| Location) | 0 | 0 | 4.47 | 2.12 | 169.1 |
| | Gradient * Size + (1 \| Location) | 0.01 | 0.12 | 3.21 | 1.79 | 156.3 |
| **$SES_{MPD}$** | (1 \| Location) | 0 | 0 | 1.03 | 1.01 | 111.7 |
| | Gradient + (1 \| Location) | 0 | 0 | 0.93 | 0.96 | 107.8 |
| | Size + (1 \| Location) | 0 | 0 | 0.96 | 0.98 | 109.1 |
| | Gradient + Size + (1 \| Location) | 0 | 0 | 0.86 | 0.93 | 104.9 |
| | Gradient * Size + (1 \| Location) | 0 | 0 | 0.61 | 0.78 | 91.5 |

**Table S2**. Parameter estimates of the best-fit candidate models ($\Delta_i$ <2) explaining the effects of Interior (I) – Edge (E) – Matrix (M) gradient of continuous forest (CF) and different-sized fragments (F100 = 100 ha, F10 = 10 ha, and F1 = 1 ha) on phylogenetic richness (PD and $SES_{PD}$) and divergence (MPD and $SES_{MPD}$). Estimates with $P < 0.05$ are marked in bold.

| **Model** | | | **Coefficient** | **Std. Error** | **t value** | **Pr (> \|t\|)** |
|---|---|---|---|---|---|---|
| **PD ~ gradient + size** | Intercept | | 351.75 | 15.36 | 22.90 | **<0.001** |
| | gradient | I | Reference | | | |
| | | E | -105.50 | 20.95 | -5.04 | **<0.001** |
| | | M | -98.88 | 20.95 | -4.72 | **<0.001** |
| | size | CF | Reference | | | |
| | | F100 | -13.72 | 25.94 | -0.53 | 0.60 |
| | | F10 | 2.99 | 22.76 | 0.13 | 0.90 |
| | | F1 | -117.69 | 22.76 | -5.17 | **<0.001** |
| **$SES_{PD}$ ~ gradient** | Intercept | | -0.99 | 0.23 | -4.40 | **<0.001** |
| | Gradient | I | Reference | | | |
| | | E | -0.42 | 0.36 | -1.16 | 0.25 |
| | | M | -1.05 | 0.36 | -2.93 | **0.006** |
| **MPD ~ gradient** | Intercept | | 52.20 | 0.59 | 88.94 | **<0.001** |
| | gradient | I | Reference | | | |
| | | E | -0.48 | 0.94 | -0.51 | 0.61 |
| | | M | -2.15 | 0.94 | -2.29 | **0.03** |
| **MPD ~ null** | Intercept | | 51.46 | 0.41 | 127.20 | **<0.001** |
| **MPD ~ size** | Intercept | | 52.14 | 0.62 | 84.34 | **<0.001** |
| | size | CF | Reference | | | |
| | | F100 | -1.81 | 1.16 | -1.57 | 0.13 |
| | | F10 | 0.35 | 1.01 | 0.35 | 0.73 |
| | | F1 | -2.09 | 1.01 | -2.07 | **0.04** |
| **MPD ~ gradient + size** | Intercept | | 52.58 | 0.67 | 78.65 | **<0.001** |
| | gradient | I | Reference | | | |
| | | E | -0.27 | 0.91 | -0.30 | 0.77 |
| | | M | -1.94 | 0.91 | -2.13 | **0.04** |
| | size | CF | Reference | | | |
| | | F100 | -1.52 | 1.13 | -1.34 | 0.19 |

| | | | | | | |
|---|---|---|---|---|---|---|
| | | F10 | 0.65 | 0.99 | 0.66 | 0.52 |
| | | F1 | -1.79 | 0.99 | -1.81 | 0.08 |
| **$SES_{MPD}$ ~ null** | Intercept | | -1.38 | 0.16 | -8.41 | **<0.001** |
| **$SES_{MPD}$ ~ gradient** | Intercept | | -1.19 | 0.24 | -4.90 | **<0.001** |
| | gradient | I | Reference | | | |
| | | E | 0.02 | 0.39 | 0.04 | 0.97 |
| | | M | -0.69 | 0.39 | -1.77 | 0.09 |

**Table S3**. Results of multiple pairwise comparisons of linear models testing for differences in phylogenetic richness (Faith's PD) across the Interior - Edge - Matrix gradient of continuous forest (CF) and different-sized fragments (1 ha, 10 ha, and 100 ha). Significant (adjusted P < 0.05) differences are highlighted in bold.

| Assemblages | Estimate | Std. Error | t value | Pr(>\|t\|) |
|---|---|---|---|---|
| CF edge - CF interior | -90.637 | 32.588 | -2.781 | 0.237 |
| CF matrix - CF interior | -111.918 | 32.588 | -3.434 | 0.065 |
| 100 ha interior - CF interior | 32.323 | 38.212 | 0.846 | 0.999 |
| 100 ha edge - CF interior | -145.078 | 38.212 | -3.797 | **0.028** |
| 100 ha matrix - CF interior | -131.676 | 38.212 | -3.446 | 0.063 |
| 10 ha interior - CF interior | 28.380 | 32.588 | 0.871 | 0.999 |
| 10 ha edge - CF interior | -137.590 | 32.588 | -4.222 | **0.010** |
| 10 ha matrix - CF interior | -85.103 | 32.588 | -2.612 | 0.314 |
| 1 ha interior - CF interior | -171.701 | 32.588 | -5.269 | **0.001** |
| 1 ha edge - CF interior | -184.389 | 32.588 | -5.658 | **0.000** |
| 1 ha matrix - CF interior | -200.257 | 32.588 | -6.145 | **0.000** |
| CF matrix - CF edge | -21.281 | 39.912 | -0.533 | 1.000 |
| 100 ha interior - CF edge | 122.959 | 44.622 | 2.756 | 0.247 |
| 100 ha edge - CF edge | -54.441 | 44.622 | -1.220 | 0.981 |
| 100 ha matrix - CF edge | -41.039 | 44.622 | -0.920 | 0.998 |
| 10 ha interior - CF edge | 119.017 | 39.912 | 2.982 | 0.164 |
| 10 ha edge - CF edge | -46.953 | 39.912 | -1.176 | 0.986 |
| 10 ha matrix - CF edge | 5.534 | 39.912 | 0.139 | 1.000 |
| 1 ha interior - CF edge | -81.064 | 39.912 | -2.031 | 0.662 |
| 1 ha edge - CF edge | -93.752 | 39.912 | -2.349 | 0.461 |
| 1 ha matrix - CF edge | -109.620 | 39.912 | -2.747 | 0.251 |
| 100 ha interior - CF matrix | 144.241 | 44.622 | 3.232 | 0.100 |
| 100 ha edge - CF matrix | -33.160 | 44.622 | -0.743 | 1.000 |
| 100 ha matrix - CF matrix | -19.758 | 44.622 | -0.443 | 1.000 |
| 10 ha interior - CF matrix | 140.298 | 39.912 | 3.515 | 0.054 |
| 10 ha edge - CF matrix | -25.672 | 39.912 | -0.643 | 1.000 |
| 10 ha matrix - CF matrix | 26.815 | 39.912 | 0.672 | 1.000 |
| 1 ha interior - CF matrix | -59.783 | 39.912 | -1.498 | 0.926 |
| 1 ha edge - CF matrix | -72.471 | 39.912 | -1.816 | 0.789 |
| 1 ha matrix - CF matrix | -88.339 | 39.912 | -2.213 | 0.547 |
| 100 ha edge - 100 ha interior | -177.401 | 48.881 | -3.629 | **0.042** |
| 100 ha matrix - 100 ha interior | -163.998 | 48.881 | -3.355 | 0.078 |
| 10 ha interior - 100 ha interior | -3.943 | 44.622 | -0.088 | 1.000 |
| 10 ha edge - 100 ha interior | -169.912 | 44.622 | -3.808 | **0.028** |
| 10 ha matrix - 100 ha interior | -117.426 | 44.622 | -2.632 | 0.304 |
| 1 ha interior - 100 ha interior | -204.023 | 44.622 | -4.572 | **0.004** |
| 1 ha edge - 100 ha interior | -216.712 | 44.622 | -4.857 | **0.002** |
| 1 ha matrix - 100 ha interior | -232.579 | 44.622 | -5.212 | **0.001** |
| 100 ha matrix - 100 ha edge | 13.402 | 48.881 | 0.274 | 1.000 |

| | | | | |
|---|---|---|---|---|
| 10 ha interior - 100 ha edge | 173.458 | 44.622 | 3.887 | **0.023** |
| 10 ha edge - 100 ha edge | 7.489 | 44.622 | 0.168 | 1.000 |
| 10 ha matrix - 100 ha edge | 59.975 | 44.622 | 1.344 | 0.963 |
| 1 ha interior - 100 ha edge | -26.622 | 44.622 | -0.597 | 1.000 |
| 1 ha edge - 100 ha edge | -39.311 | 44.622 | -0.881 | 0.999 |
| 1 ha matrix - 100 ha edge | -55.178 | 44.622 | -1.237 | 0.979 |
| 10 ha interior - 100 ha matrix | 160.056 | 44.622 | 3.587 | **0.046** |
| 10 ha edge - 100 ha matrix | -5.914 | 44.622 | -0.133 | 1.000 |
| 10 ha matrix - 100 ha matrix | 46.573 | 44.622 | 1.044 | 0.994 |
| 1 ha interior - 100 ha matrix | -40.025 | 44.622 | -0.897 | 0.998 |
| 1 ha edge - 100 ha matrix | -52.713 | 44.622 | -1.181 | 0.985 |
| 1 ha matrix - 100 ha matrix | -68.581 | 44.622 | -1.537 | 0.913 |
| 10 ha edge - 10 ha interior | -165.970 | 39.912 | -4.158 | **0.012** |
| 10 ha matrix - 10 ha interior | -113.483 | 39.912 | -2.843 | 0.211 |
| 1 ha interior - 10 ha interior | -200.080 | 39.912 | -5.013 | **0.001** |
| 1 ha edge - 10 ha interior | -212.769 | 39.912 | -5.331 | **0.001** |
| 1 ha matrix - 10 ha interior | -228.636 | 39.912 | -5.729 | **0.000** |
| 10 ha matrix - 10 ha edge | 52.487 | 39.912 | 1.315 | 0.968 |
| 1 ha interior - 10 ha edge | -34.111 | 39.912 | -0.855 | 0.999 |
| 1 ha edge - 10 ha edge | -46.799 | 39.912 | -1.173 | 0.986 |
| 1 ha matrix - 10 ha edge | -62.667 | 39.912 | -1.570 | 0.902 |
| 1 ha interior - 10 ha matrix | -86.598 | 39.912 | -2.170 | 0.575 |
| 1 ha edge - 10 ha matrix | -99.286 | 39.912 | -2.488 | 0.380 |
| 1 ha matrix - 10 ha matrix | -115.154 | 39.912 | -2.885 | 0.196 |
| 1 ha edge - 1 ha interior | -12.688 | 39.912 | -0.318 | 1.000 |
| 1 ha matrix - 1 ha interior | -28.556 | 39.912 | -0.715 | 1.000 |
| 1 ha matrix - 1 ha edge | -15.868 | 39.912 | -0.398 | 1.000 |

**Table S4**. Results of simple Mantel tests between beta diversity metrics (Pβ and COMDIST) and geographic distance between sites at the Biological Dynamics of Forest Fragments Project, Brazil. Pearson's product-moment correlation and Spearman's rank correlation were used to test for linear and non-linear relationships, respectively.

| **Metric** | | **Pearson's product-moment correlation** | | **Spearman's rank correlation rho** | |
|---|---|---|---|---|---|
| | | **r** | ***P*** | **r** | ***P*** |
| Pβ | $\beta_{total}$ | **0.134** | **0.013** | **0.117** | **0.024** |
| | $\beta_{repl}$ | 0.031 | 0.299 | 0.024 | 0.339 |
| | $\beta_{rich}$ | 0.059 | 0.138 | 0.049 | 0.162 |
| COMDIST | | 0.055 | 0.230 | 0.021 | 0.368 |

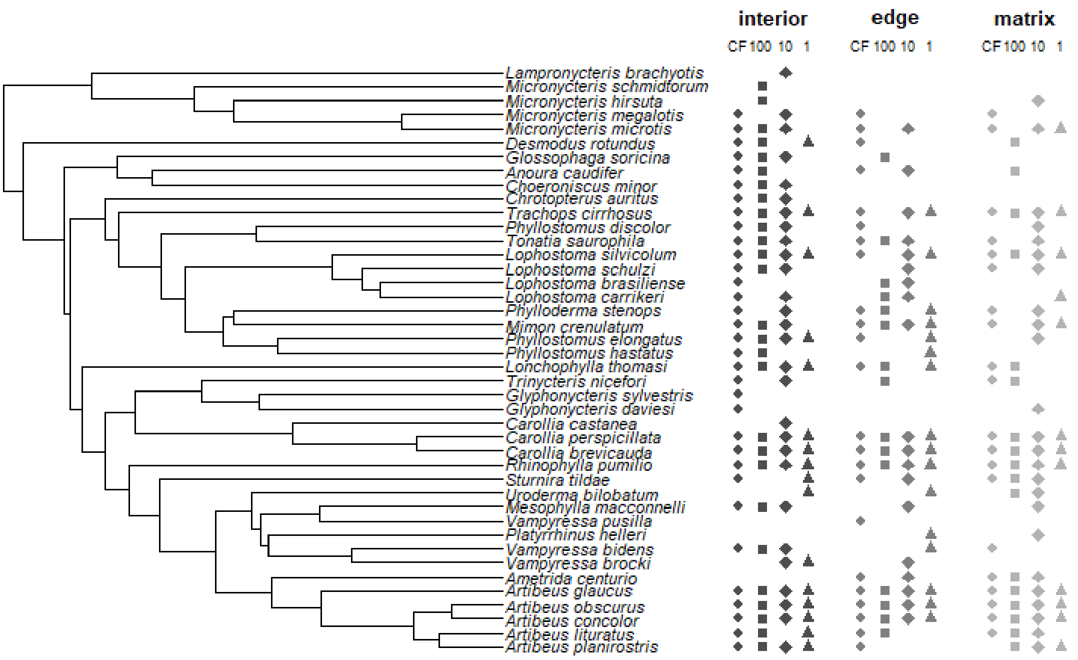


**Fig. S1** ~~The p~~Phylogenetic relationship of phyllostomid bats at the BDFFP and their occurrence in continuous forest (CF, circle), 100 ha fragments (square), 10 ha fragments (diamond) and 1 ha fragments (triangle) across a gradient of interior, edge, and matrix sites.
